# Isolation-free Identification and Phenotyping of First-Trimester Extravillous Trophoblasts Residing in Cervical Fluid

**DOI:** 10.1101/2025.09.02.673885

**Authors:** Manpreet Kalkat, Kevin D. Pavelko, Michael Strausbach, Austin Goodyke, Lauren C. Crimp, Dennis Lee, D. Randall Armant, Sascha Drewlo

## Abstract

Preeclampsia (PE) remains difficult to predict, particularly when it manifests late in gestation. To capture early placental signals, we profiled trophoblast cells sampled from the cervix in the first trimester using mass cytometry (CyTOF). We established protocols for clinical sample storage and applied spike-in reference control cells to deliver reproducible, batch-corrected protein measurements, thereby advancing CyTOF from a discovery tool to a translational platform. Within HLA-G⁺CD45⁻ cells, we identified canonical CK7⁺ extravillous trophoblasts, as well as a previously unrecognized CK7⁻subset, and both subsets expressed placental proteins. Expression of PAPP-A, GAL-13, and GAL-14 was significantly altered in a pilot cohort of pregnancies that subsequently developed late-onset PE, distinguishing cases from controls at both single-marker and multivariate levels. These findings reveal unexpected trophoblast heterogeneity, demonstrate that placental alterations are detectable before the development of late-onset PE, and establish cervical trophoblast profiling as a promising platform for scalable biomarker discovery and first-trimester risk assessment in placenta-mediated disorders.

**Impact Statement:** First-trimester trophoblasts sampled from the cervix reveal early molecular changes associated with late-onset preeclampsia, while an isolation-free, reference-normalized CyTOF workflow establishes a scalable, clinically compatible platform for biomarker discovery and multicenter-ready early risk assessment in pregnancy.

## Introduction

Pregnancy complications that arise from disordered placental development, collectively termed malplacentation, remain a leading cause of maternal and fetal morbidity. Examples include preeclampsia (PE) and fetal growth restriction (FGR), which both stem from aberrant trophoblast invasion and incomplete spiral artery remodeling during the first trimester (Brosens, Robertson, and Dixon 1970; Pijnenborg et al. 1983; Roberts and Lain 2002; Lindheimer, Roberts, and Cunningham 2009; Roberts and Escudero 2012). Interventions, such as low dose aspirin (Bujold, Roberge, and Nicolaides 2014) or Pravastatin (Ramma and Ahmed 2014), when administered before 16 weeks, have shown potential in mitigating PE’s occurrence among high-risk patients; however, the efficacy is contingent upon dispensation during the pivotal vascular remodeling phase in the first trimester. The placenta is effectively inaccessible during this developmental period, making early diagnosis difficult and delaying intervention until pathology is well-established and possibly irreversible. Current screening relies on maternal serum markers such as Placental Growth Factor (PlGF) and Soluble fms-like Tyrosine Kinase-1 (sFlt-1) together with Doppler ultrasound (Bart et al. 2025). Unfortunately, the predictive power of these tools peaks only in the late second trimester, well beyond the optimal window for preventive therapy.

PE itself is phenotypically heterogeneous (Magee, Nicolaides, and von Dadelszen 2022; Redman, Staff, and Roberts 2022). Early-onset disease, diagnosed before 34 weeks, is tightly linked to overt placental pathology marked by shallow trophoblast invasion, hypoxic villi, and pronounced endothelial dysfunction. Late-onset PE, which manifests at or after 34 weeks, presents a more nuanced picture. Some investigators frame it primarily as a maternal cardiovascular syndrome precipitated by the metabolic demands of late pregnancy (Valensise et al. 2008; Roberts and Hubel 2009), whereas others argue that subtler, cumulative placental stress beginning early in development eventually triggers disease (Chaiworapongsa et al. 2014; Burton et al. 2019; Redman, Staff, and Roberts 2022). Malplacentation due to trophoblast dysfunction, depending on its severity and timing, could contribute to a broad spectrum of pregnancy disorders, including miscarriage, FGR, PE and various associated pathologies (Burton and Jauniaux 2004). These observations suggest that detailed interrogation of trophoblast biology across all trimesters could ultimately provide insights into susceptibility across the entire spectrum of PE and FGR.

Trophoblast Retrieval and Isolation from the Cervix (TRIC) first demonstrated that extravillous trophoblast (EVT) cells, distinguished by their expression of Human Leukocyte Antigen G (HLA-G) (Chumbley et al. 1994; McMaster et al. 1995), are naturally shed into the endocervical canal and can be harvested for analysis non-invasively early in gestation (Bolnick et al. 2014; Jain et al. 2016; Moser et al. 2018). Immunocytochemical labeling of the reproductive tract during early pregnancy with an antibody to HLA-G reveals EVT cells migrating from the decidua into the uterine cavity where they can be carried by secreted mucus to the cervix (Moser et al. 2018). The protein expression patterns of EVT cells obtained from the cervix (cEVT) by TRIC and examined by immunocytochemistry are significantly altered during the first trimester in pregnancies that later succumb to early pregnancy failure (Fritz, Kohan-Ghadr, Bolnick, et al. 2015) or develop complications (PE or FGR) in the third trimester (Bolnick et al. 2016), suggesting the diagnostic potential of cEVT cells for malplacentation. However, TRIC depends on manual manipulation of cell populations, an expertise-intensive step that constrains throughput and standardization. To overcome this limitation, we developed an isolation-free alternative that couples cervical fluid collection by the Pap procedure with high-dimensional mass cytometry (CyTOF) (Muftuoglu and Andreeff 2024; Spitzer and Nolan 2016). The workflow preserves sample integrity from collection to analysis, incorporates spiked-in reference cell lines for cross-run normalization, and uses a curated antibody panel to resolve cEVT populations within heterogeneous cervical specimens. By eliminating manual cEVT isolation, this scalable and reproducible platform delivers non-invasive, first-trimester access to placental biology and pathology, bridging a critical diagnostic gap for both early– and late-onset PE, as well as other disorders of malplacentation.

## Methods

### Patient selection

The institutional review board of Corewell Health approved this study, and each participating patient provided written informed consent. For this study, pregnant women (*n* = 25) between 10 – 14 weeks of gestational were recruited at Corewell Health at 221 Michigan OB/GYN, ICCB/Beltline OB/GYN, 3800 Lake Michigan, and the OB/GYN Residency Clinics. Pregnant mothers, with no active infections at time of collection (i.e. herpes, HIV, Hepatitis C, sexually transmitted infections) scheduled for a pelvic exam or clinical cervical swab, and deemed safe to swab (no cerclage, dilation, active bleeding) were consented to specimen collection protocol with clinical data abstraction under Corewell Health IRB protocol CHW 2017-198. Patient data was deidentified and abstracted directly into the study’s REDCap database. Following consent, a sterile cervical swab was used to gently swab the vaginal walls, rotating for 15-30 seconds, to obtain secretions from the mucosal membrane and immediately transferred into cold DMEM:F12 media (Gibco), as previously described (Imudia et al. 2009).

### Clinical sample processing

Mucosal collections were transferred to the Grand Rapids Research Center and processed within 4 hours of collection. In brief, mucosal collections were gently dissociated by pipetting and cells were collected via centrifugation. Cells were then resuspended into PBS (Gibco) and filtered through a 100µM filter. Cells were again collected by centrifugation and fixed in 2% paraformaldehyde (EMS) in PBS for 15 minutes with gentle agitation. Finally, cells were resuspended in PBS with 0.1% BSA (Sigma), 0.5% sodium azide (Thermo) and 0.04% paraformaldehyde and stored at 4C until shipment to the Mayo Clinic for CyTOF analysis.

### Cell lines

Untransduced cell lines were obtained from American Type Culture Collection (ATCC, Manassas, Virginia, USA) including Chinese Hamster Ovary (CHO-K1, #CCL-61), Human Embryonic Kidney (HEK-293, #CRL-1573), human Primary Peripheral Blood Mononuclear (PBMC, #PCS-800-11), and two human placenta-derived trophoblast (JEG-3 and HTR-8/SVneo, #HTB-36 and #CRL-3271, respectively) cells. For HLA-G over-expressing cells, an HLA-G-encoding lentiviral expression construct (Cat# RC205216L4V) was obtained from Origene Technologies Inc. (Rockville, MD, USA) and transduced into HEK-293 cells, following manufacturer protocols. The remaining overexpressing cells were made from lentiviral expression constructs obtained from Genecopoeia (Rockville, MD, USA) and transduced into CHO-K1 cells as per manufacturer protocols, including constructs encoding Pregnancy Associated Pregnancy Protein A (PAPP-A) (Cat# LPP-Y4134-Lv213-050), PlGF (Cat# LPP-CS-Z0166-Lv213-01-050), alpha-fetoprotein (AFP) (Cat# LPP-G0171-Lv207-050), Flt1 (Cat# LPP-CS-K4570-Lv213-01-050) or empty control Lv207 (Cat# LPP-NEG-Lv207-050) and Lv213 (Cat# LPP-NEG-Lv213-050). Cells lines were cultured under recommended conditions, which were DMEM for CHO-K1 and HEK-293 cells, RPMI 1640 for PBMC, JEG-3 and HTR-8/SVneo cells, all supplemented with 10% FBS and 1% penicillin-streptomycin. For transduction, cells were seeded at ∼50% confluence one day prior to infection and then incubated with lentiviral particles at a multiplicity of infection (MOI) between 5–10, in the presence of 8 μg/mL polybrene (Sigma-Aldrich) to enhance viral uptake. After 24 hours, viral supernatant was replaced with fresh complete medium. At 48 hours post-transduction, cells were subjected to selection using puromycin or hygromycin at the optimal killing concentration (determined by a prior kill curve assay for each cell line). Selection was continued for 5–7 days until all non-transduced control cells were eliminated. Individual clones were isolated by flow sorting of reporter fluorescent proteins (EGFP or mCherry) using a BD FACSAria Fusion Flow Cytometer (BD Biosciences, Franklin Lakes, NJ, USA). Stable cell lines were expanded and maintained under continuous puromycin or hygromycin selection.

Reference cell lines were generated to verify antibody specificity (**Supplemental Figure S1**). Upon verification, the reference panel was reduced to minimize the number of required barcodes for experimental efficiency (**Table 1**). A representative gating strategy and associated protein expression profiles of the EVT-specific antibody panel are shown in **Supplemental Figure S2 and S3**. For this study, we operationally define EVT cells as HLA-G⁺/CD45⁻ cells detected in cervical samples, and subsequent use of the term ‘trophoblast’ refers specifically to this EVT population. Use of CD45 to negatively screen cells eliminates known expression of HLA-G acquired by certain immune cells (Amodio et al. 2013; Hsu et al. 2012; Rouas-Freiss et al. 2021).

**Table 1.** Trophoblast Panel – Antibody Marker References / Controls.

| No. | Cell Line | CD105/<br>Endoglin | Ck7 | VEGFR1/<br>FLT1 | PAPP-A | PLGF | CD45 | AFP | GAL-13 | EpCAM | GAL-14 | HLA-G | Casp3 |
| --- | --- | --- | --- | --- | --- | --- | --- | --- | --- | --- | --- | --- | --- |
| 1 | LV207-CHO-eGFP | - | - | - | - | - | - | - | + | - | + | - | +/- |
| 2 | LV213-CHO-mCherry | - | - | - | - | - | - | - | + | - | + | - | +/- |
| 3 | JEG3 | + | + | - | +/- | +/- | - | - | + | + | + | +/- | +/- |
| 4 | HTR8 | - | - | - | - | - | - | - | + | - | + | - | +/- |
| 5 | PBMC | - | - | - | - | - | + | - | - | - | - | - | +/- |
| 6 | CHO-PAPP-A-mCherry | - | - | - | + | - | - | - | + | - | + | - | +/- |
| 7 | CHO-PLGF G4 | - | - | - | - | + | - | - | + | - | + | - | +/- |
| 8 | CHO-AFP-eGFP | - | - | - | - | - | - | + | + | - | + | - | +/- |
| 9 | HLA-G-HEK293 (EE) | - | - | - | - | - | - | - | + | - | + | + | +/- |
| 10 | CHO-Flt1 | - | - | + | - | - | - | - | + | - | + | - | +/- |

### Preparation of Barcoded Control Cell Lines for CyTOF

Reference cell lines (**Table 1**) were thawed and counted. After thawing, each cell line was incubated with cisplatin solution for live/dead discrimination. Cells were washed with cell staining buffer and PBS. After washing, cells were fixed with 2% paraformaldehyde for 20 minutes with gentle agitation, washed, and then resuspended in Maxpar® Barcode Perm Buffer (Cat. No. 201057, Standard BioTools). Maxpar® Palladium Barcoding Kit (Cat. No. 201060, Standard BioTools), barcodes 1 through 10, were added to each tube of cells and incubated for 15 minutes at room temperature.

Cells were washed, and all reference cell lines were then combined into a single tube. Aliquots of 1 million total cells were cryopreserved in 10% DMSO in complete media (10% FBS/RPMI).

### CyTOF Sample Processing

Fixed trophoblast samples were removed from 4°C storage, or cryopreserved samples were thawed prior to staining. Cells were pelleted, washed and counted. One to 3 million cells were aliquoted into individual Eppendorf tubes. Cells were resuspended in barcoding permeabilization buffer prior to addition of barcoding reagent corresponding to palladium barcodes 11–20 in the Maxpar Cell-ID 20-Plex Pd Barcoding Kit (Cat. No. 201060, Standard BioTools). After barcoding, all experimental samples and an aliquot of the barcoded reference cell lines were pooled into a single tube. Combined cells were pelleted and washed with cell staining buffer. A cocktail of antibodies to the extracellular markers (**Table 2**) was added to the resuspended cells and incubated at room temperature with gentle agitation for 45 minutes. After cell surface staining cells were washed, fixed with 2% paraformaldehyde and then permeabilized and resuspended in PERM-S solution. A cocktail of antibodies to the intracellular markers was then added to the cells and incubated at 4°C for 45 minutes. Cells were washed and then resuspended in Cell-ID™ Intercalator-Ir (Cat. No. 201192A, Standard BioTools) for 30 minutes or overnight at 4 °C. Intercalated samples were then washed with cell staining buffer and PBS prior to resuspension in a 1:10 mixture of EQ Four Element Calibration Beads and cell acquisition solution. The sample was then filtered through a 70 µm blue cap Falcon tube and then loaded onto the Helios mass cytometer sample loader for data acquisition. The Helios mass cytometer was tuned according to the manufacturer’s recommendations prior to use and all events were collected at 100-200 events per second.

**Table 2.**
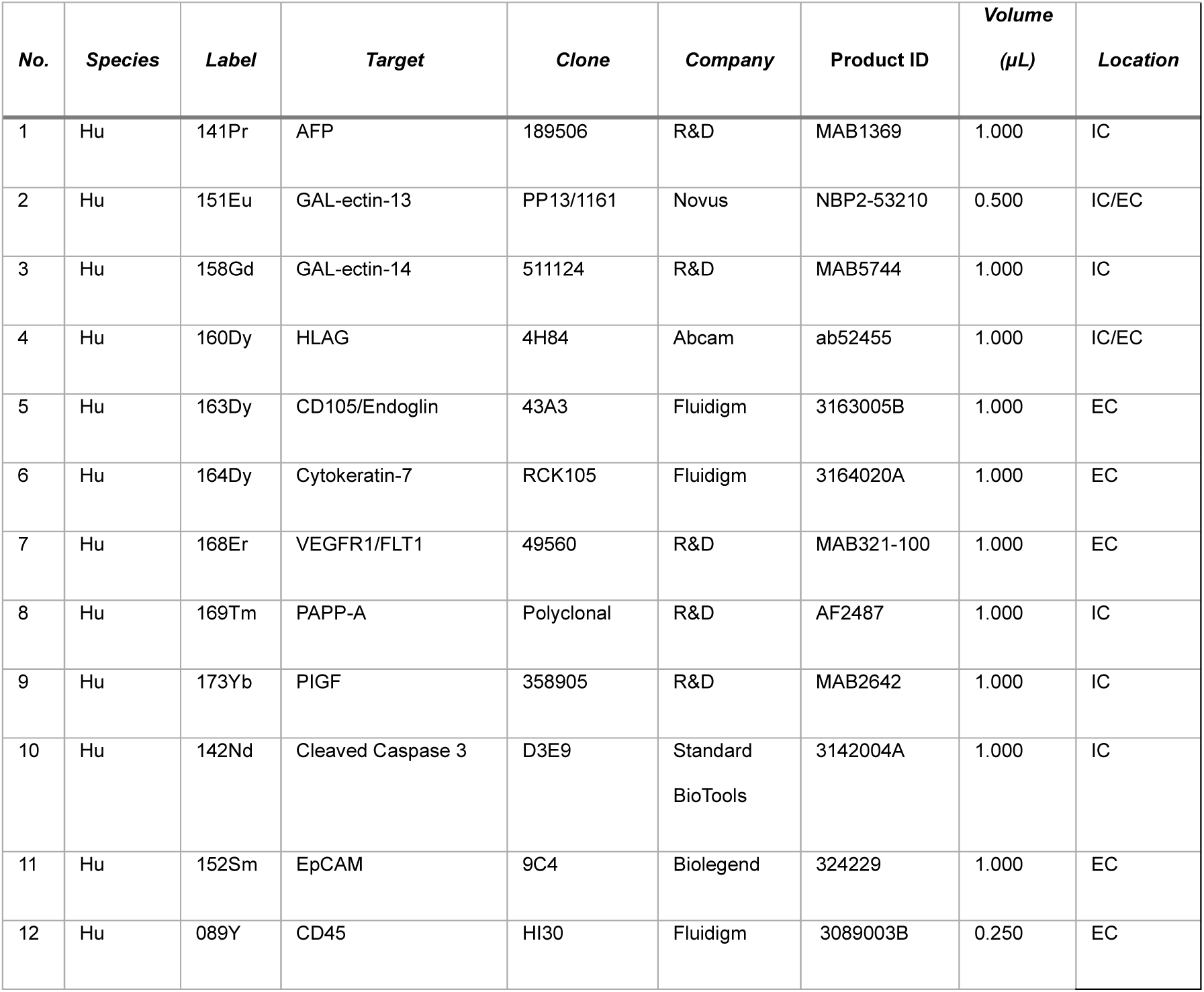
Trophoblast Antibody Panel.

| **No.** | **Species** | **Label** | **Target** | **Clone** | **Company** | **Product ID** | **Volume**<br>**(μL)** | **Location** |
| --- | --- | --- | --- | --- | --- | --- | --- | --- |
| 1 | Hu | 141Pr | AFP | 189506 | R&D | MAB1369 | 1.000 | IC |
| 2 | Hu | 151Eu | GAL-ectin-13 | PP13/1161 | Novus | NBP2-53210 | 0.500 | IC/EC |
| 3 | Hu | 158Gd | GAL-ectin-14 | 511124 | R&D | MAB5744 | 1.000 | IC |
| 4 | Hu | 160Dy | HLA-G | 4H84 | Abcam | ab52455 | 1.000 | IC/EC |
| 5 | Hu | 163Dy | CD105/Endoglin | 43A3 | Fluidigm | 3163005B | 1.000 | EC |
| 6 | Hu | 164Dy | Cytokeratin-7 | RCK105 | Fluidigm | 3164020A | 1.000 | EC |
| 7 | Hu | 168Er | VEGFR1/FLT1 | 49560 | R&D | MAB321-100 | 1.000 | EC |
| 8 | Hu | 169Tm | PAPP-A | Polyclonal | R&D | AF2487 | 1.000 | IC |
| 9 | Hu | 173Yb | PIGF | 358905 | R&D | MAB2642 | 1.000 | IC |
| 10 | Hu | 142Nd | Cleaved Caspase 3 | D3E9 | Standard<br>BioTools | 3142004A | 1.000 | IC |
| 11 | Hu | 152Sm | EpCAM | 9C4 | Biolegend | 324229 | 1.000 | EC |
| 12 | Hu | 089Y | CD45 | HI30 | Fluidigm | 3089003B | 0.250 | EC |

### CyTOF Data Analysis

After acquisition, all FCS files were normalized and debarcoded using CyTOF Software v7.0.8493 (Standard BioTools). Batch normalization to reference cell lines was performed using the CytoNorm plug-in v1.0 in FlowJo. Data were cleaned using FlowJo v10.10.0 (BD Biosciences) or OMIQ v3.7.0 (OMIQ, Inc.) following the gating strategies outlined in **Supplemental Figure S2 and S3**. Additional data analysis, including clustering, was performed in either FlowJo or OMIQ.

## Results

Endocervical swabs were leveraged as a minimally invasive source of trophoblast cells, which were processed and analyzed by CyTOF to generate high-dimensional placental protein profiles. The study workflow (**Figure 1**) summarizes the overall approach, from patient sampling through standardized analysis, and represents the first application of CyTOF for analysis of cEVT cells. A central goal of the workflow design is to support larger multi-center clinical studies with the ability to store and normalize patient samples obtained at different times. Building on this foundation, we evaluated sample stability, defined trophoblast heterogeneity, and explored associations between placental protein signatures and pregnancy outcomes. For these analyses, we developed a 12-marker CyTOF panel that captured placental identity and disease markers [Alpha Fetoprotein [alpha-fetoprotein (AFP), Pregnancy-associated Plasma Protein A (PAPP-A), Galectin-13 (GAL-13), Galectin-14 (GAL-14), and human leukocyte antigen G (HLA-G)], epithelial differentiation [Cytokeratin 7 (CK7), Epithelial Cell Adhesion Molecule (EpCAM)], angiogenic regulation [(Endoglin (CD105)] and apoptosis (Cleaved Caspase-3), while excluding immune contaminants (CD45). This analyte panel formed the foundation of the study workflow illustrated in **Figure 1** and was applied throughout subsequent experiments. We additionally validated antibody specificity using overexpressing reference cell lines (**Supplemental figure S1**) and then reduced the reference panel to a minimal set for all further experiments (**Table 1 and Supplemental figure S3**).

**Figure 1.**
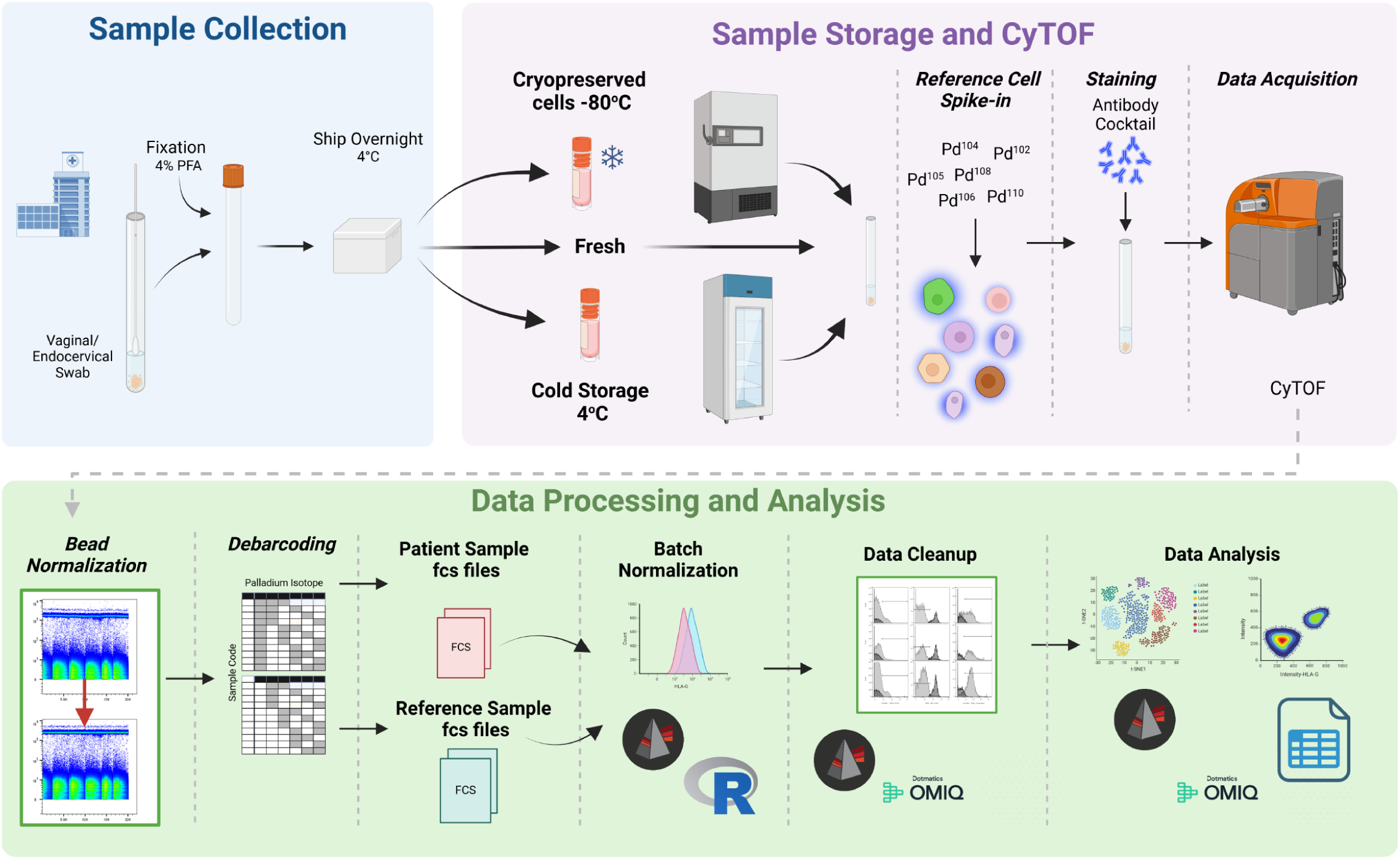
Sample collection, storage and CyTOF workflow for evaluation of endocervical swab samples. A coordinated system was established to facilitate sample movement through the CyTOF workflow. Cells derived from endocervical swabs were collected at Corewell Health, Grand Rapids, MI, and were fixed with 4% paraformaldehyde to ensure sample consistency over time and stabilize antigen epitopes. Samples were then shipped overnight at 4°C. For stability tests, samples were analyzed within 2 or 3 days as fresh samples or were stored for 30 days at 4°C or cryopreserved at –80°C. Preserved reference samples were spiked into the clinical test samples and analyzed by CyTOF. Data files were normalized to spiked-in EQ beads to stabilize mass isotope signals and batch normalization was performed using spiked in reference cells when necessary. All data analysis was performed using the cytometry analysis programs FlowJo and/or OMIQ.

We used CyTOF to profile patient-derived endocervical specimens and define the molecular composition of trophoblast populations (**Figure 2**). Four individual endocervical samples were processed by CyTOF and analyzed using an established gating strategy (**Supplemental figures S2 and S3**). Dimensionality reduction by UMAP revealed clear separation of trophoblasts from other cell types, characterized by strong expression of HLA-G and absence of CD45 **(Figure 2 and Supplemental Figure S4)**. Within the HLA-G–positive population, two discrete regions were consistently identified, suggesting heterogeneity among endocervical trophoblasts. These regions were manually gated region 1 and 2 for downstream analysis and compared across a panel of phenotyping markers. Both regions expressed proteins characteristic of placental origin, including PAPP-A, GAL-13, and GAL-14, to confirm their trophoblastic identity.

**Figure 2.**
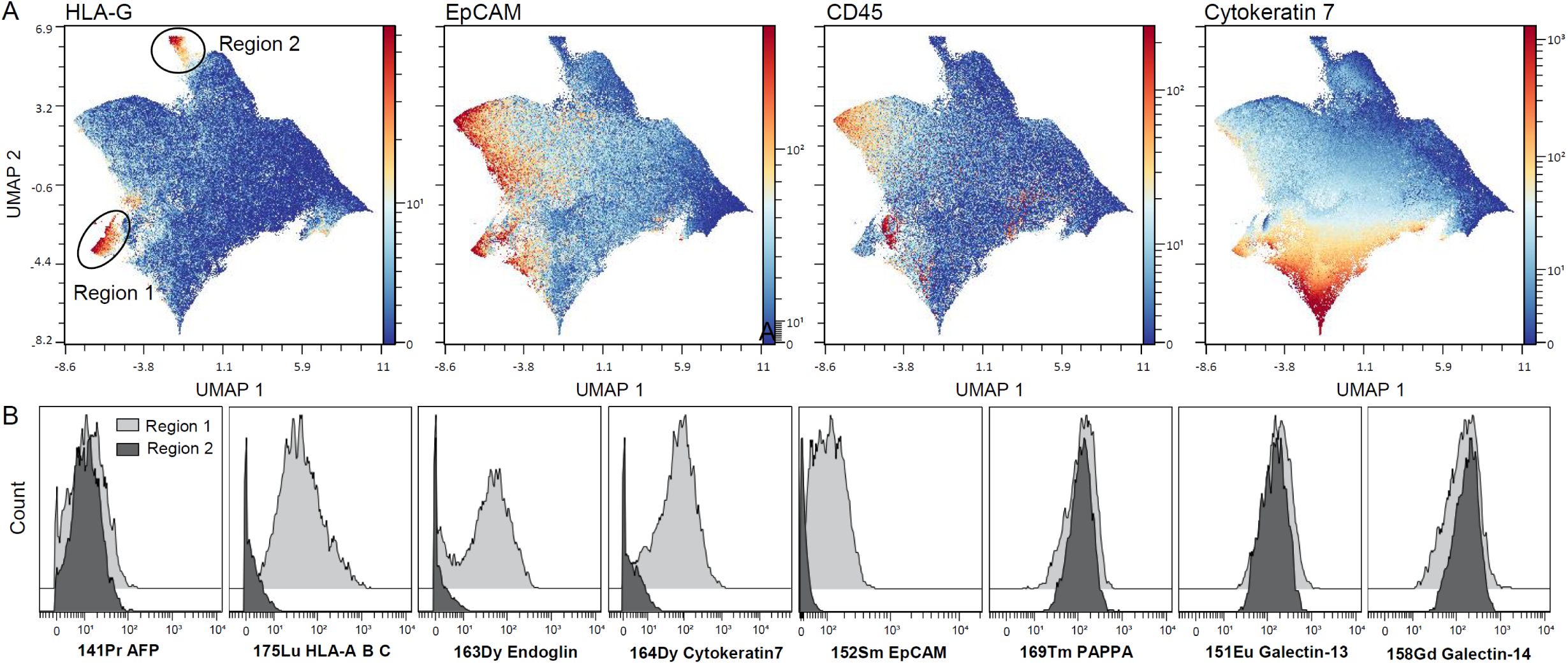
Evaluation of patient samples by CyTOF. Four independent cell samples derived from endocervical swabs were analyzed by CyTOF. Pooled data that were processed using an established gating strategy were analyzed using the dimensionality reduction technique UMAP. **A)** UMAP projections with overlying expression densities for HLA-G, EpCAM, CD45 and CK7. Two unique regions of HLA-G expression were manually gated for downstream phenotyping. **B)** Expression levels of 7 phenotyping markers were measured in the two HLA-G expressing regions identified in A. It identified differential expression levels in region 1 compared to region 2.

However, the two regions diverged in their expression of EpCAM, CK7, Endoglin, and HLA-ABC, suggesting molecular differences that distinguish these two subsets, with region 2 cells being negative for these 4 markers. Region 1 cells clearly have an EVT molecular phenotype, while the Region 2 cells may constitute an alternative EVT subtype or a different cell type altogether. Together, these findings reveal that endocervical HLA-G+/CD45– cells are not homogeneous, but instead comprise at least two phenotypically distinct cervical fluid subpopulations detectable by CyTOF.

Building on this observation of heterogeneity within the HLA-G+ population, we next employed CK7 expression as a distinguishing feature to further resolve two cellular subpopulations. We first gated on HLA-G+/CD45– cells from representative endocervical swab samples to exclude immune cells **(Figure 3).** Within this population, two subsets were distinguished based on CK7 expression. CK7+ cells demonstrated uniform expression of six placental markers—AFP, GAL-13, GAL-14, PAPP-A, EpCAM, and Endoglin—consistent with a differentiated EVT phenotype. In contrast, CK7– cells expressed AFP, GAL-13, GAL-14, and PAPP-A, but showed little to no expression of EpCAM or Endoglin, indicating a more restricted molecular profile. These findings reveal that endocervical HLA-G+ non-immune cells comprise at least two subsets defined by CK7 status, with CK7+ cells exhibiting a broader placental EVT signature compared to CK7– cells.

**Figure 3.**
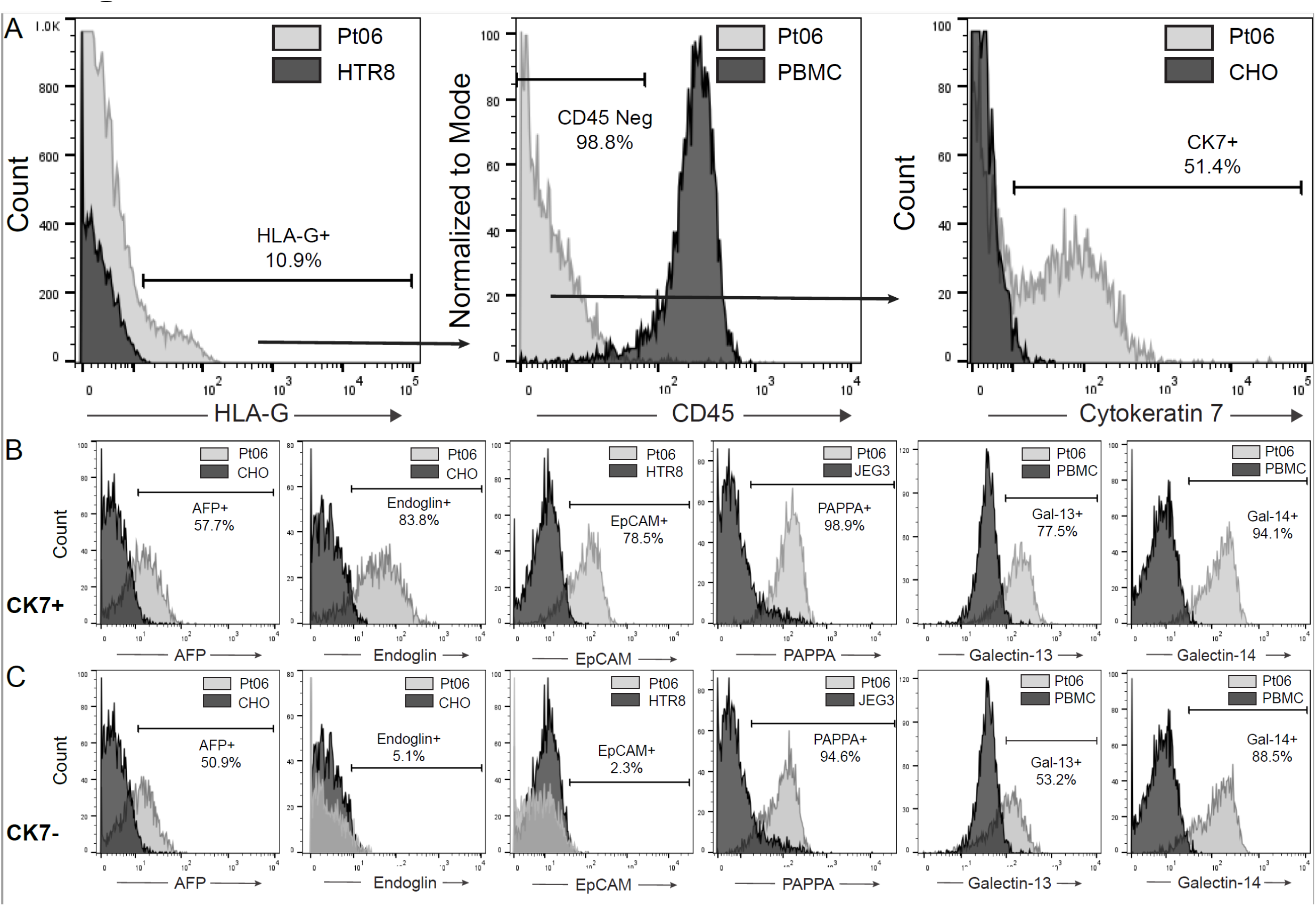
Identification of unique populations of HLA-G+ cells differentiated by expression of cytokeratin 7 (CK7). A representative endocervical cell sample from patient 06 was evaluated by CyTOF. **A)** Gating strategy using spike in reference cells to define gates. Trophoblast cells were identified by HLA-G+ and CD45-cell populations. HLAG+ CD45-cells were identified to have both CK7+ and CK7-cell populations (right panel). **B)** Expression of 6 phenotyping markers in the CK7+ cells. C) Expression the same 6 phenotyping markers in the CK– cells. Percentages for all histograms indicate the percentage positive relative to established negative control cell lines.

Having defined CK7+ and CK7– EVT-like subsets of cells, we next evaluated whether these populations could be reliably detected after extended sample storage, a critical step for enabling batched clinical studies **(Figure 4).** Endocervical swab-derived cells were tested immediately or preserved for 27 days at either –80°C or 4°C and then analyzed by CyTOF. In representative samples, expression of HLA-G, CD45, and CK7 remained stable across both conditions, indicating preservation of key lineage-defining markers. Quantification across four individual patient samples confirmed that the relative abundance of HLA-G+ cells, the proportion of CD45– trophoblasts, and the frequency of CK7+ cells were comparable between fresh and stored samples. These findings demonstrate that extended storage, whether by refrigeration or cryopreservation, does not compromise the capacity to identify trophoblast populations, enabling patient samples to be reliably analyzed in batches, independently of when they arrive in the laboratory.

**Figure 4.**
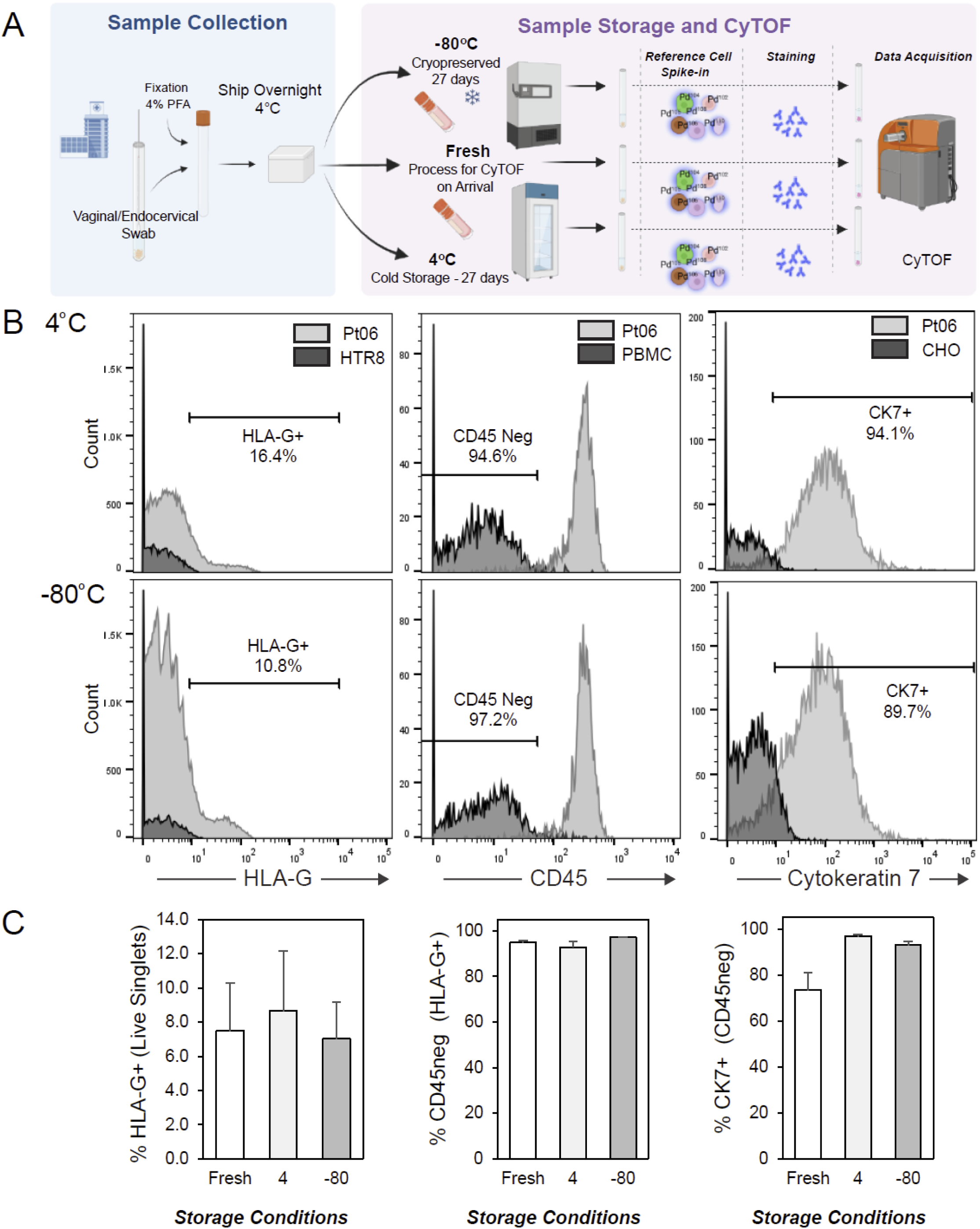
Effect of storage conditions on the outcome of trophoblast characterization by CyTOF. **A)** Schema for testing the effect of storage at –80°C or 4°C on samples stored for 27 days. **B)** Stability of HLA-G, CD45 and CK7 expression in samples stored for 27 days. Representative sample derived from patient 06 in Figure 3. **C)** Comparison of the relative percentage of HLA-G+, CD45⁻ and CK7+ cells across subjects (n=4 patient samples).

Having established reliable storage conditions and a normalization strategy with spike-in reference cells, we next applied the workflow to a small (n=21; 17 controls, 4 late onset-PE) cohort study (**Table 3 and Figure 5**). Endocervical swabs were stored at –80°C until batch processing by CyTOF, with preserved reference cells included in each run to ensure consistent normalization across experiments. This approach enabled direct comparison of control pregnancies and those with pathological outcomes. Differential expression analysis revealed that PAPP-A, GAL-13, and GAL-14 were significantly altered between groups, with all three markers reduced in the pathology (PE) cohort. These findings demonstrate that, once storage and normalization workflows were optimized, CyTOF could reproducibly identify placental protein signatures in early pregnancy samples that aligned with later adverse outcomes.

**Figure 5.**
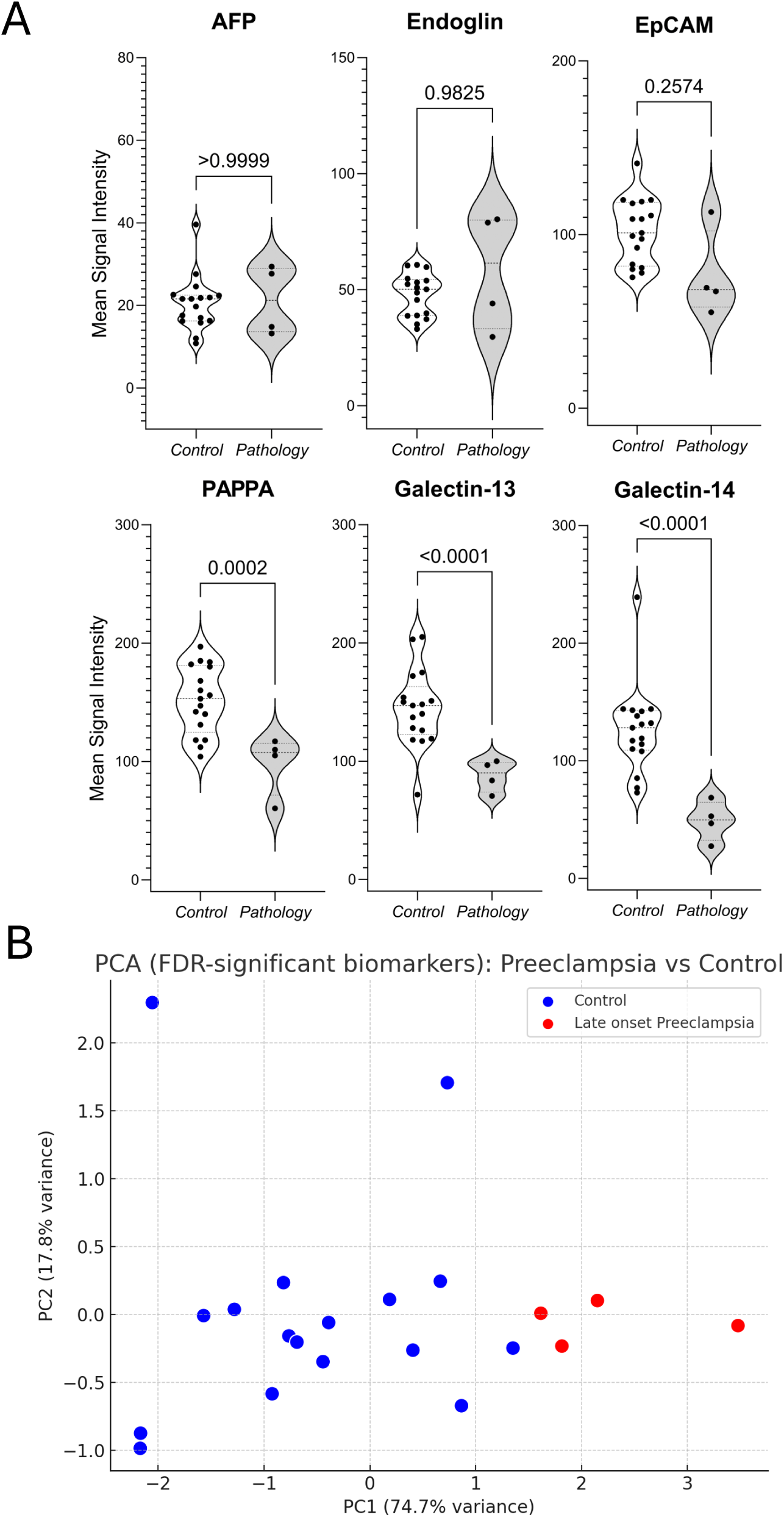
Differential expression of trophoblast CyTOF markers aligns with pregnancy outcomes. **A)** Endocervical samples were evaluated by CyTOF from control pregnancies and pregnancies with pathologic sequelae resulting in PE. Expression of PAPP-A, GAL-13 and GAL-14 differed significantly between control and pathology groups (T-test). **B)** Principal component analysis (PCA) of patient samples based on only the biomarkers that differed significantly (GAL-14, PAPP-A, and GAL-13). Each point represents one patient, plotted along the first two principal components. Blue circles represent control samples, while red circles represent pathology, which was late-onset PE in all cases. The separation of groups indicates that these biomarkers capture key variance distinguishing PE from controls.

**Table 3.** Patient Characteristics.

| Characteristic | Control | Late onset Preeclampsia |
| --- | --- | --- |
| N | 17 | 4 |
| Maternal Age (years) | 30.4 ± 3.6 | 31.5 ± 0.6 |
| BMI | 24.7 ± 4.9 | 29.4 ± 6.9 |
| Gestational Age at Delivery<br>(weeks) | 38.8 ± 1.2 | 37.2 ± 2.1 |
| Birthweight (g) | 3551 ± 496 | 3460 ± 1171 |
| Ethnicity: Asian | 1 (5.9%) | 0 (0.0%) |
| Ethnicity: Hispanic | 1 (5.9%) | 0 (0.0%) |
| Ethnicity: Other | 0 (0.0%) | 1 (25.0%) |
| Ethnicity: Unknown | 1 (5.9%) | 0 (0.0%) |
| Ethnicity: White | 14 (82.4%) | 3 (75.0%) |

## Discussion

This study demonstrates that first-trimester EVT cells obtained from cervical mucus (cEVT) can be profiled by CyTOF to uncover both unexpected heterogeneity and early-stage protein profiles that differentiate pregnancies destined to develop late-onset PE. Using HLA-G to identify EVT cells and CD45 to gate out maternal immune cells, two trophoblast-like populations were identified: a canonical CK7⁺ subset and an unanticipated CK7⁻ subset that nevertheless expressed the hallmark placental proteins PAPP-A, GAL-13, and GAL-14. These findings expand the current definition of EVT identity and establish cervical sampling as a feasible, minimally invasive approach for studying placental biology during pregnancy.

The discovery of a CK7⁻ trophoblast population adds further depth to this picture. While CK7 has long been considered a defining marker of the epithelial trophoblast lineage, our findings reveal that some HLA-G+/CD45– cells lack CK7 yet maintain robust expression of known secreted trophoblast proteins. The consistent detection of PAPP-A and placental Galectins in this subset raises the possibility that the CK7⁻ cells could have specialized roles in trophoblast secretory or immunomodulatory function. Functional studies will be required to define these roles.

The most significant finding is that PAPP-A, GAL-13, and GAL-14 expression was altered significantly in first trimester EVT cells from pregnancies that subsequently developed late-onset PE. These changes distinguished cases from controls both at the level of individual marker distributions and through principal component analysis, where late-onset PE samples clustered separately. This is striking, as late-onset PE has often been regarded as less tightly linked to placental dysfunction, as compared to early-onset disease. Our data suggest that even in late-onset PE, early placental alterations are measurable months before clinical presentation.

Beyond biological discovery, this study also introduces a methodological framework that will support future clinical translation. We established a protocol for long-term storage of cervical samples compatible with CyTOF, enabling flexible biobanking and retrospective analysis. In addition, we developed a normalization strategy using spike-in reference controls to adjust for batch effects and address inter-run drift, improving reproducibility. Together, these advances provide a technical foundation for applying high-dimensional single-cell proteomics to large pregnancy cohorts, a critical step toward clinical implementation.

The use of spike-in reference controls makes this approach scalable beyond a single laboratory setting. By anchoring protein expression signatures to stable external references, inter-run and inter-site variability can be minimized, allowing data generated across different centers and time points to be directly compared. This feature is essential for validating candidate biomarkers in diverse populations and will facilitate collaborative, multi-center studies aimed at establishing standardized assays for clinical investigation. In this way, our approach not only uncovers biological signals but also addresses one of the major barriers to translating single-cell proteomics into real-world clinical testing.

From a translational perspective, the ability to capture first-trimester trophoblast molecular signatures associated with perinatal disease, specifically late-onset PE, has important clinical implications. Current screening strategies for PE rely on maternal serum biomarkers and uterine artery Doppler, which are imperfect and primarily report risk for early-onset disease. Our findings suggest that cEVT analysis could provide an orthogonal, cell-based window into placental function, complementing existing tools such as serum biomarkers and Doppler indices, and potentially enabling earlier identification of patients at risk for late-onset PE and other placenta-derived disorders.

This study has limitations. The cohort sizes were modest, which limits power to detect subtler differences beyond PAPP-A and placental Galectins. Adverse outcomes were limited to a single disorder, late-onset PE. However, these observations extend and align with our prior TRIC studies encompassing 41 pregnancies analyzed by immunohistochemistry, which also identified dysregulated placental protein signatures before disease onset (Bolnick et al. 2016). Together, these two studies support the plausibility of early placental contributions to perinatal disease. Because cEVT cells were not purified, maternal cellular admixture is possible, mitigated by HLA-G⁺/CD45⁻ gating. Mechanistic studies are still needed to define how reduced PAPP-A and placental Galectin expression shapes EVT behavior and contributes to disease.

Importantly, the reduced expression of biomarkers aligns with established placental biology. PAPP-A increases local Insulin-like Growth Factor (IGF) bioavailability by cleaving Insulin-like Growth Factor Binding Protein 4 (IGFBP-4, supporting EVT invasion and arteriolar remodeling), and low first-trimester PAPP-A levels are associated with placenta-mediated complications, including PE and FGR (Oxvig 2015). GAL-13 is linked to PE risk in serum meta-analyses, and GAL-14 promotes EVT migration/invasion via Akt–MMP9/N-cadherin signaling, and decreased placental expression of GAL-13/14 has been reported in PE (Than et al. 2014; Wu, Liu, and Ding 2021; Wang et al. 2021). These findings are further supported in studies using TRIC to isolate cEVT cells early in gestation and assess biomarker dysregulation (Bolnick et al. 2014; Fritz, Kohan-Ghadr, Sacher, et al. 2015). Consistent with the hypothesis that both early– and late-onset PE develop downstream of syncytiotrophoblast stress (Redman, Staff, and Roberts 2022), the identification of first-trimester molecular changes facilitated by our storage and reference-normalization workflow provides a translational proof-of-concept platform for multicenter validation.

In summary, we report that CyTOF interrogation of cervical mucus reveals unexpected CK7 heterogeneity of cEVT cells and demonstrates that alterations in PAPP-A, GAL-13, and GAL-14 expression are detectable in the first trimester of pregnancies that subsequently develop late-onset PE. By establishing protocols for clinical sample storage and introducing a reproducible normalization strategy, we provide both biological insight and a technical framework for future biomarker discovery and clinical studies. These findings challenge the prevailing view that late-onset PE is largely a maternal disorder by showing that early placental contributions are detectable *in vivo* during the first trimester. In contrast to syncytiotrophoblast stress models derived from delivered placentas, cervical EVT profiling provides a unique early-pregnancy window and establishes isolation-free, reference-normalized CyTOF as a multicenter-ready platform for clinical translation across placenta-mediated disorders.

## Figure Legends

**Supplemental Figure S1.**
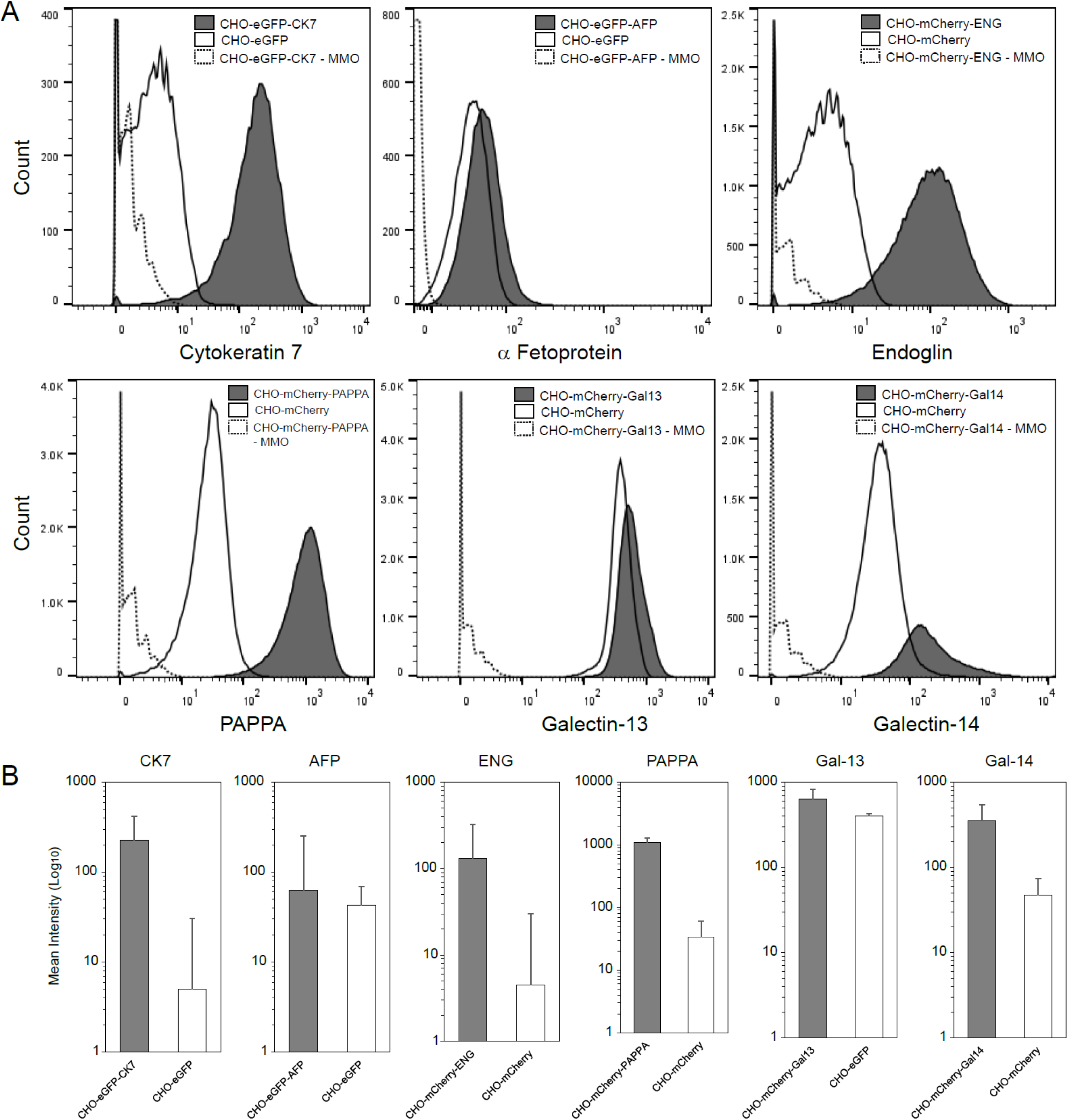
CyTOF expression in cell lines generated to overexpress key components of the trophoblast reference panel. A. Expression of CK7, AFP, Endoglin/CD105, PAPP-A, GAL-13 and GAL-14 in overexpressing cell lines. MMO – metal minus one condition to control for signal spillover. B. Mean signal intensity of control and overexpressing cell lines for the above targets.

**Supplemental Figure S2.**
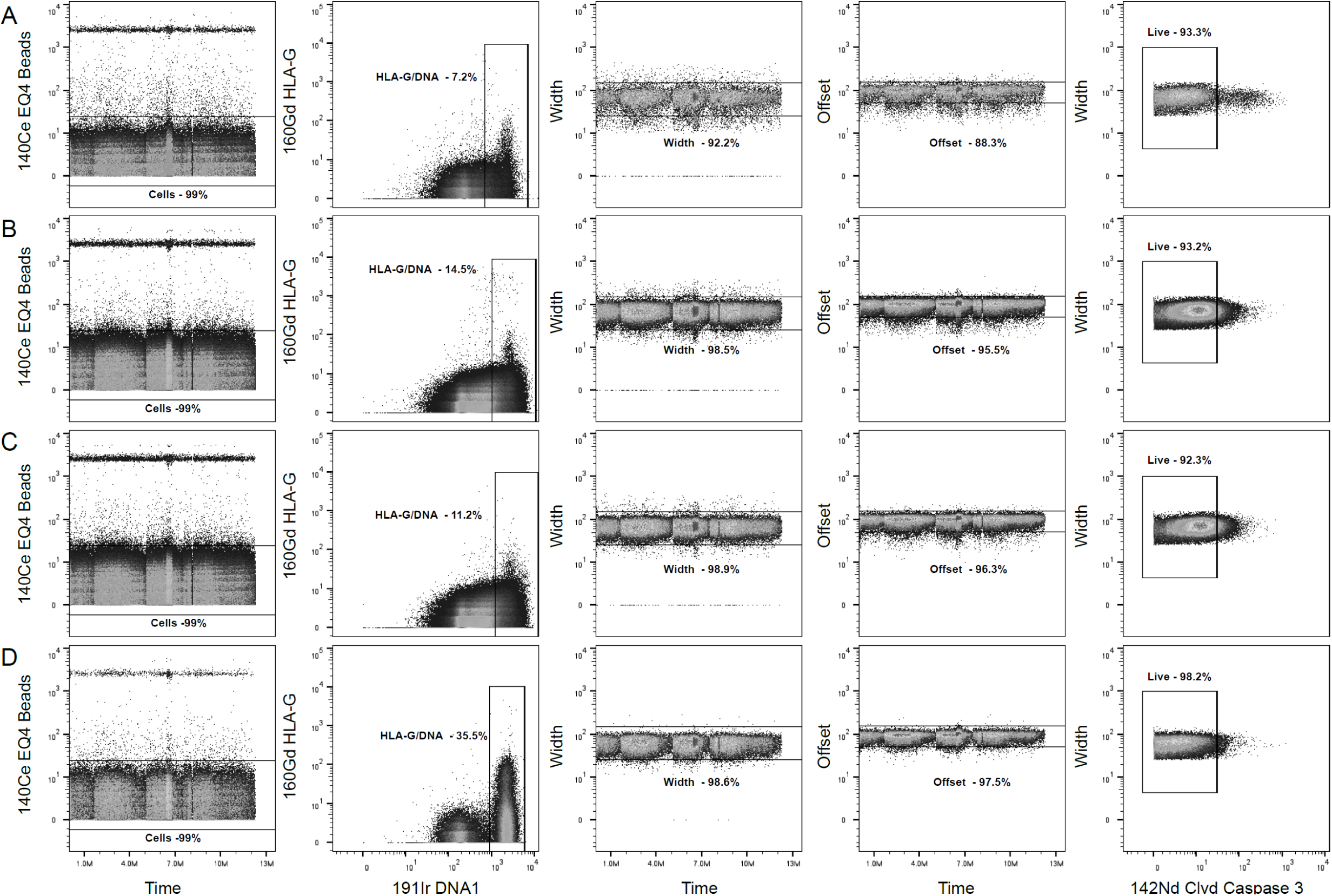
CyTOF sample cleanup gating strategy. Four independent samples were gated on EQ beads, DNA content, width, offset and live cells.

**Supplemental Figure S3.**
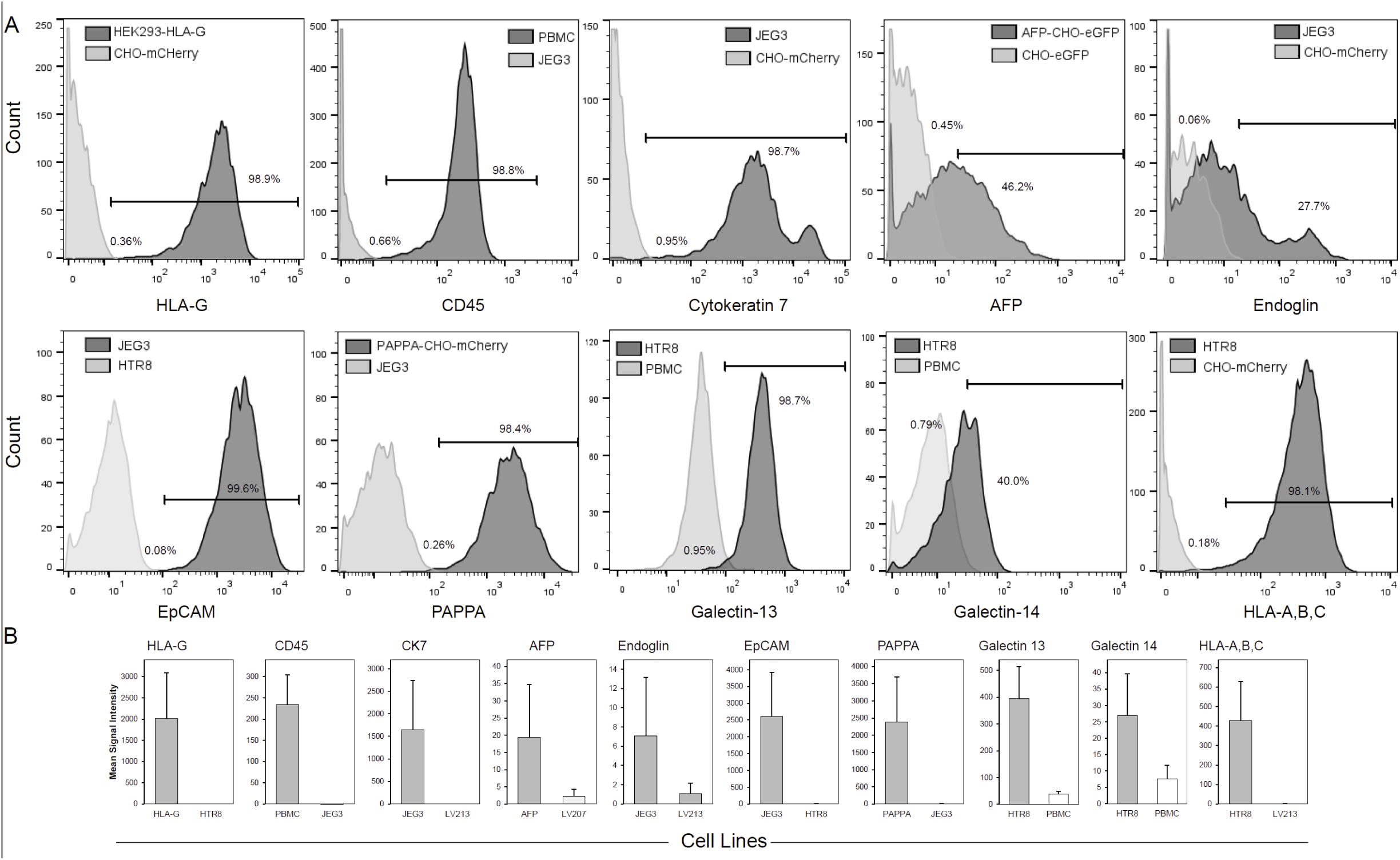
Gating strategy based on a minimal reference spike-in cell line panel, used to define positive and negative thresholds for cell population identification. A. Cells overexpressing the target of interest were used to define cutoff values for positive and negative expression profiles. For CD45, peripheral blood mononuclear cells (PBMCs) were used to determine CD45 positivity, while an epithelial cell line, JEG3 cells, were used to define the negative population. JEG3 cells were used to define EpCAM positive gating. To reduce the number of required spike ins, HTR8 cells were used to define GAL-13, GAL-14 and HLA-ABC positivity, and served as a negative control for EpCAM expression. B. Mean signal intensity for positive and negative controls for the trophoblast antibody panel.

**Supplemental Figure S4.**
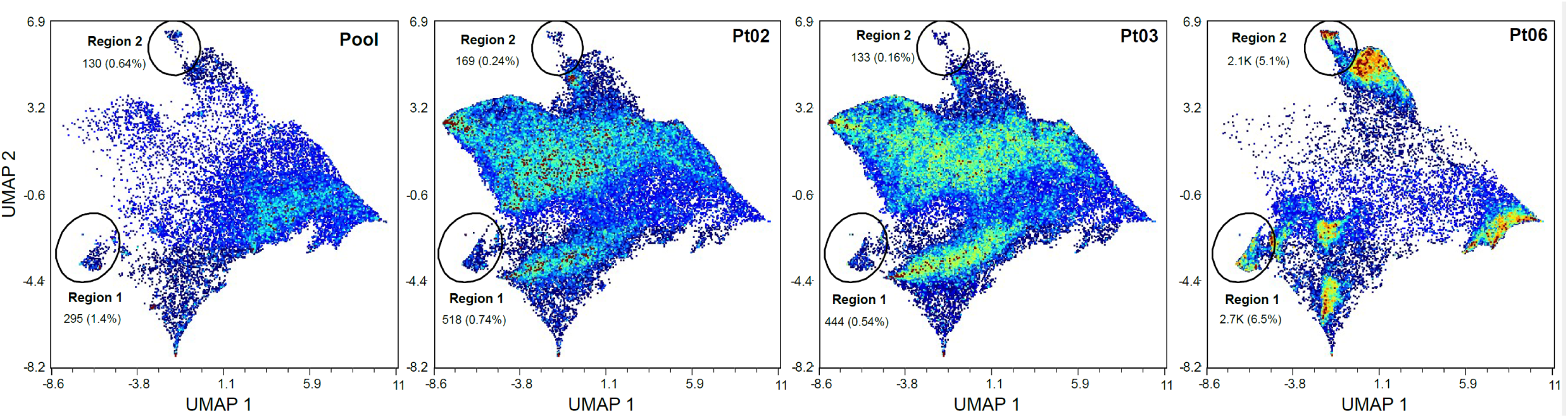
Evaluation of four individual patient samples by CyTOF. Four independent cell samples derived from endocervical swabs were analyzed by CyTOF. UMAP projections for individual patients are colored by cell density.

